# MIST-Explorer: The Comprehensive Toolkit for Spatial Omic Analysis and Visualization of Single-Cell MIST Array Data

**DOI:** 10.1101/2025.04.29.650640

**Authors:** Clark Fischer, Jianxin Chen, Arafat Meah, Jun Wang

## Abstract

Recent advances in spatial proteomics have enabled high-dimensional protein analysis within tissue samples, yet few methods accurately detect low-abundance functional proteins. Spatial MIST (Multiplex *In Situ* Tagging) is one such technique, capable of profiling over 100 protein markers spatially at single-cell resolution on tissue sections and cultured cells. However, despite the availability of various open-source tools for image registration and visualization, no dedicated software exists to align the images and analyze spatial MIST data effectively. To address this gap, we present MIST-Explorer, a comprehensive, user-friendly toolkit for the visualization and analysis of single-cell spatial MIST array data. Developed in Python with a PyQt6-based graphical interface, MIST-Explorer streamlines the spatial omics workflow—from image preprocessing and registration to cell segmentation and protein quantification. The software supports two workflows: one for preprocessed datasets and another for raw image inputs, ensuring broad compatibility across experimental designs. Key features include tile-based image registration using Astroalign and PyStackReg, deep learning-based segmentation with StarDist, multi-channel visualization with layer controls, and an interactive analysis module offering ROI selection along with histograms, heatmaps, and UMAP plots. MIST-Explorer generates spatially resolved expression tables readily compatible with downstream single-cell analysis pipelines. By integrating all major steps into a single platform, MIST-Explorer empowers researchers to derive biological insights from complex spatial omics datasets without requiring extensive computational expertise.

**Availability and implementation:** Freely available at https://github.com/MIST-Explorer/MIST-Explorer.

## 1 Introduction

Recent advances in single-cell proteomics have allowed researchers to analyze hundreds to thousands of proteins with high spatial resolution on tissue and cell samples. However, these methods often require special, expensive instrumentation that most labs do not possess, or they are time-consuming that needs days to process one sample. In addition, the mass spectrometry based single-cell proteomics methods cannot detect many functional proteins, which are often in low abundance and are vital to understanding disease mechanisms. Spatial multiplex *in situ* tagging (MIST) utilizes a microbead array that enables the “printing” of tissue molecules onto the array. Spatial information is preserved in this process and a sequential fluorescence technique is used to decode the identity of the protein. This method has been shown to detect over 100 protein markers with high sensitivity (Meah et al., 2023).

In this paper, we highlight the MIST-Explorer, a custom platform designed to assist data processing of MIST experimental results. With the MIST-Explorer, typical processes including registration, visualization, and downstream analysis are all carried out since they have been integrated into a single, user-friendly GUI application. Our tool has the potential to be applied to many other spatial proteomics and transcriptomics applications such as those described in Chen et al., 2020. These applications require precise alignment of fluorescence images to preserve spatial information. Our tool outputs coordinates of each cell’s center along with expression values — a format readily compatible with single-cell level spatial transcriptomics analysis. Coupled with the spatial MIST and the derived technologies, MIST-Explorer can significantly accelerate biomedical research, drug discovery and clinical studies.

## 2 Methods

### 2.1 Preparation of DNA-Conjugated Microbeads

Amine-functionalized polystyrene beads (2 µm; Life Sciences Technologies) were conjugated with two distinct single-stranded DNA (ssDNA) sequences per bead, each corresponding to a specific protein target (sequences listed in Table S1). Beads were activated with 10 mM BS3 (Thermo Fisher) for 20 min, centrifuged, and washed. They were then incubated with 0.05% Poly-L-lysine (PLL; Ted Pella) for 2 hours to enhance DNA attachment, followed by reaction with 5 mM Azido-PEG4-NHS (Vector Laboratories) for 4 hours under agitation. Separately, 150 µM ssDNAs were conjugated with DBCO-NHS (Click Chemistry) at 5 mM, pH 8.5, for 4 hours. Modified oligonucleotides were purified using 7 kDa MWCO Zeba spin columns and incubated overnight with functionalized beads to achieve dual-oligo loading. Beads were washed with Milli-Q water, resuspended, and assembled into a microarray on glass slides. Monolayer formation and uniformity were confirmed microscopically.

### 2.2 Preparation and Purification of DNA-Conjugated Antibodies

Antibodies were conjugated to biotinylated and sequence k’-tagged oligos using DBCO-azido click chemistry. Antibodies were concentrated to 1 mg/mL and modified with UV-cleavable azido-NHS ester (1:15 molar ratio, 2 hours). In parallel, oligos (200 µM, pH 8.0) were conjugated with UV-cleavable DBCO-NHS ester (1:20 molar ratio, 2 hours). Both were purified with 7 kDa MWCO Zeba columns, mixed, and incubated overnight. Conjugates were further purified via FPLC and stored at 0.3–0.5 mg/mL in PBS with 0.05% NaN□ at 4°C.

### 2.3 Tissue Handling and Spatial MIST Clamping

Formalin-fixed paraffin-embedded (FFPE) kidney tissues were deparaffinized in xylene, rehydrated in graded ethanol, rinsed in water, and subjected to antigen retrieval (citrate buffer, pH 6.0, 95°C, 20 min). Sections were permeabilized with 0.3% Triton X-100 and blocked with 5% goat serum, salmon sperm DNA, and Tween 20. Tissues were incubated with UV-cleavable, oligo-conjugated antibodies (biotin and k′-tagged) for 3 hours at RT, followed by nuclear staining with POPO-1 (1:200, 15 min). MIST arrays were blocked, PEG-treated, and clamped to tissue sections. UV exposure (365 nm, 5 min) transferred oligos and dye to the array. Arrays were washed and incubated sequentially with streptavidin, biotinylated antibody, complementary k-DIG oligo, anti-DIG antibody, and fluor-labeled secondary antibodies, enabling dual-protein visualization.

### 2.4 Imaging and MIST Decoding

Arrays were imaged using an Olympus VS200 scanner under brightfield, DAPI, Cy3, and Cy5 channels. For decoding, arrays were denatured with 1 M NaOH (2 min), washed, and hybridized with fluorophore-labeled complementary DNAs in 40% formamide/10% dextran sulfate buffer for 1 hour. Two decoding cycles were performed using distinct fluor-tagged oligos. All images were computationally aligned using in-house software, enabling precise decoding of bead identities and quantification of protein targets.

### 2.5 Implementation of MIST-Explorer

MIST-Explorer was written in Python 3.14 with the PyQt6 library to create a native platform and quick application. The graphical user interface (GUI) was built using PyQT6. Core functionalities rely on established scientific Python libraries including NumPy and Pandas for data manipulation, Matplotlib for generating interactive visualizations, and scikit-image for image processing tasks. StarDist was used for deep-learning-based cell segmentation. Astroalign and PyStackReg were used for image registration purposes. Performance-critical algorithms, such as protein visualization rendering, utilize Just-in-Time (JIT) compilation via Numba. The software is open-source and available at https://github.com/MIST-Explorer/MIST-Explorer.

### 2.6 Data

MIST-Explorer supports two primary workflows for data input, accommodating users with varying levels of pre-processed data:

Workflow 1: Loading Pre-processed Segmentation Mask and Data Table: This workflow is intended for users who have performed cell segmentation and feature quantification externally. The required inputs are:

1. Cell Segmentation Mask: A single-channel image file (e.g., TIFF format) where pixels corresponding to each identified cell are assigned a unique integer ID.
2. Quantitative Data Table: A tabular data file (CSV or Excel format) containing quantified measurements (e.g., mean protein expression intensity, integrated density) for each cell. This table must be indexed or contain a column with unique cell identifiers that directly correspond to the integer IDs used in the segmentation mask image, which ensures accurate mapping between the mask and the quantitative data.

Workflow 2: Loading Raw Images for In-Application Processing: This workflow allows users to leverage the tool’s integrated preprocessing capabilities. Users can start with:

1. Target Image: A multi-layer TIFF image containing the protein and cell detection data from the microscopy experiment. Included in the TIFF image is a brightfield channel as the first layer. This serves as the target/moving image for multi-channel image alignment. Also included in the TIFF are channels for nuclear stains, cell morphology markers, and protein intensity.
2. Reference image: A multi-layer TIFF image with the first layer being the bright field of the microbead array and subsequent layers being the fluorescent intensities of varying fluorescent colors. The brightfield layer serves as the reference for multi-channel image alignment.

Users employing this workflow utilize the functions within the ‘Data Processing’ tab to perform tasks such as multi-channel image alignment and subsequent cell segmentation. This process generates the required cell segmentation mask within the application. Following segmentation, the quantitative data table (as described in Workflow 1) should be provided by the user or potentially generated through further steps to link expression values to the newly defined cell segments for visualization and analysis.

### 2.7 Image Analysis

#### 2.7.1 Data Acquisition

The single-round labeling and multi-cycle decoding technique based on the spatial MIST platform allows for the detection of hundreds of proteins simultaneously. The process involves a cocktail of UV-cleavable DNA barcoded antibodies which stain the tissue. After each staining, the DNA oligos can be liberated by UV light and be “printed” on the MIST array which comprises of microbeads conjugated with complimentary DNA sequences. Once the tissue is printed, the microarray is then scanned. This initial scan contains the nuclear marker, protein channel and Brightfield which we later use for registration purposes. Decoding the microarray entails successive cycles of DNA elution and hybridization with fluorophore□conjugated complements, with each protein defined by a unique sequence of bead color transitions based on a predetermined coding scheme (see next sections). For complete methodological details, see Meah et al. (2023).

**Figure 1.**
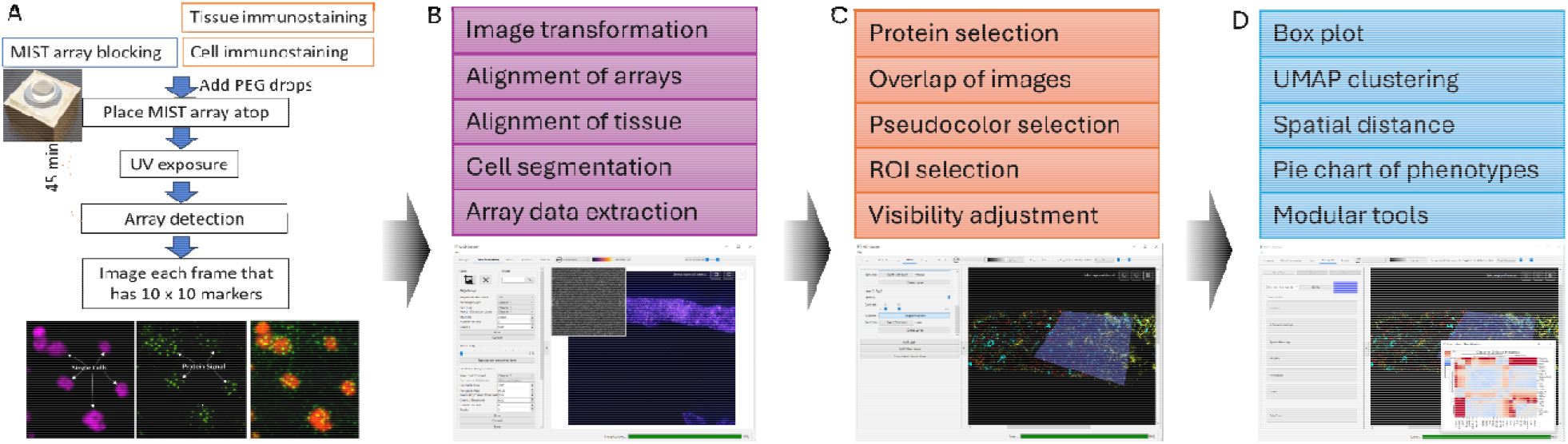
A brief overview of MIST-Explorer’s role in processing and analyzing single-cell spatial MIST data. A) Experimental flow of spatial MIST experiments on cells or tissue sections. Inset images show a fully assembled MIST array and a set of example images detecting both cell location and protein expression. B) Major modules of MIST-Explorer for data processing that includes the sequential steps. C) View tab enables visualization of data processing results. Protein expressions are shown in segmented cells. Multiple proteins can be visualized together by assigning them different pseudocolors. D) Modular bioinformatic tools can be added to analyze the spatial proteomic data in single cells. The tools are expandable by developers.

#### 2.7.2. Data Processing

MIST-Explorer includes a dedicated data processing module to prepare images for visualization and analysis. Tools are provided for interactive image cropping via user-drawn rectangular selections and image rotation by input of the desired angle of rotation. Prior to image registration, images were cropped and rotated to lower the dimensionality, resulting in increased speed and efficiency.

To accurately retrieve and visualize the expression of various proteins in cells, the target image must be aligned to the reference image. Traditional image registration algorithms involving feature detection such as Scale-invariant Feature Transform (SIFT) and its derivatives such as Speeded-Up Robust Features (SURF) rely on identifying key points distinct from their surroundings. However, since registration of the target image and the reference image depends on their respective brightfield layers, which contain millions of microbeads indistinguishable from each other, key point-based approaches yielded poor registration results. Given that the full-sized image is on the order of gigabytes, registration as-is was impractical. Thus, a tile-based approach was implemented. The images were split into smaller, equal-sized tiles with overlapping regions to ensure seamless stitching following the registration. Each tile was first registered by an affine transformation using astroalign. This library allows for the estimation of a scale, translation, and rotation invariant transform without relying on key points by calculating triangular correspondences between the fixed and moving image (Beroiz et al., 2020). This is followed by non-rigid transformation using pystackreg. For each set of fixed and moving tiles, if a transformation could not be estimated, then the transformation for the previous set of tiles was used.

In the MIST-Explorer, users can select a reference channel (optionally specifying if the blue channel, often DAPI/Hoechst, should guide initial alignment), the cell marker channel, and the protein detection channel. There are also key parameters controlling the tiled alignment, including maximum image dimension for processing (MaxSize), the number of tiles (NumberOfTiles), and the pixel overlap between tiles (Overlap), which are user-configurable to optimize alignment accuracy and performance.

After aligning the reference and target images, a Gaussian smoothing filter was applied, with adjustable intensity controlled via a slider to reduce image noise prior to subsequent analysis. The smoothed image can replace the original protein intensity layer in the target image.

Cell segmentation of the cell marker channel in the target image is required to identify the microbeads spatially. To accomplish this, StarDist was chosen for its robustness in identifying overlapping regions (Schidmt et al., 2018). The MIST-explorer integrates the StarDist deep learning model for automated cell segmentation directly within the application. Users can select the channel containing the relevant signal or load a single channel image to be segmented. Users can choose from pre-trained StarDist 2D models (e.g., 2D_versatile_fluo) and adjust critical segmentation parameters. The result is a segmentation mask image, where each pixel belonging to a specific cell carries a unique integer label.

The protein identity of each bead was determined using a pre-established color code. The fluorescence color changes from each cycle were recorded as part of the decoding process, generating a CSV containing global coordinates and color code of all microbeads. This color code is a unique identifier for a specific protein. Using cell labels and color codes, the single-cell protein expression profiles were recorded in a comprehensive CSV, containing the spatial coordinates for each cell, and the 16-bit median intensity for varying proteins within the cell. For cells with incomplete protein expression profiles (e.g., missing a bead), the median intensity of the nearest bead with the protein of interest was used to complete the profile. The nearest bead was calculated as the shortest Euclidean distance between the center of the cell and a list of bead locations with the protein of interest. The resulting comprehensive data file was used for downstream spatial protein expression visualization and bioinformatics analysis.

#### 2.7.3 Interactive Visualization Module (“View” Tab)

The core visualization interface allows for the exploration of protein expression data overlaid on microscopy images:

Layer Management: Users can dynamically add multiple layers corresponding to different protein expression channels from the input data table or import external images (PNG, JPG, TIFF) as additional layers (e.g., for annotations or reference structures).

Layer Display Controls: Each layer can be independently controlled. An opacity slider (0-100%) adjusts layer transparency for visualizing co-localization. Contrast is adjusted using dual sliders defining minimum and maximum intensity thresholds, effectively implementing percentile-based intensity windowing for enhancing visibility without altering raw data. Grayscale protein layers can be assigned user-selectable color tints using standard RGB or CMYK palettes, facilitating pseudo-colored multi-channel image composition. Individual layers can be toggled visible/invisible or deleted.

Visualization Canvas: Layers are rendered onto a central canvas using alpha compositing for smooth blending based on individual layer settings (opacity, contrast, tint).

Region of Interest (ROI) Selection: Interactive tools are provided directly on the canvas for selecting ROIs for downstream analysis. These include rectangular, circular/elliptical, and free-form polygon selection tools.

#### 2.7.4 Quantitative Analysis Module (“Analysis” Tab)

This module enables quantitative analysis of protein expression data within user-defined ROIs selected in the View tab (Figure 2):

**Figure 2.**
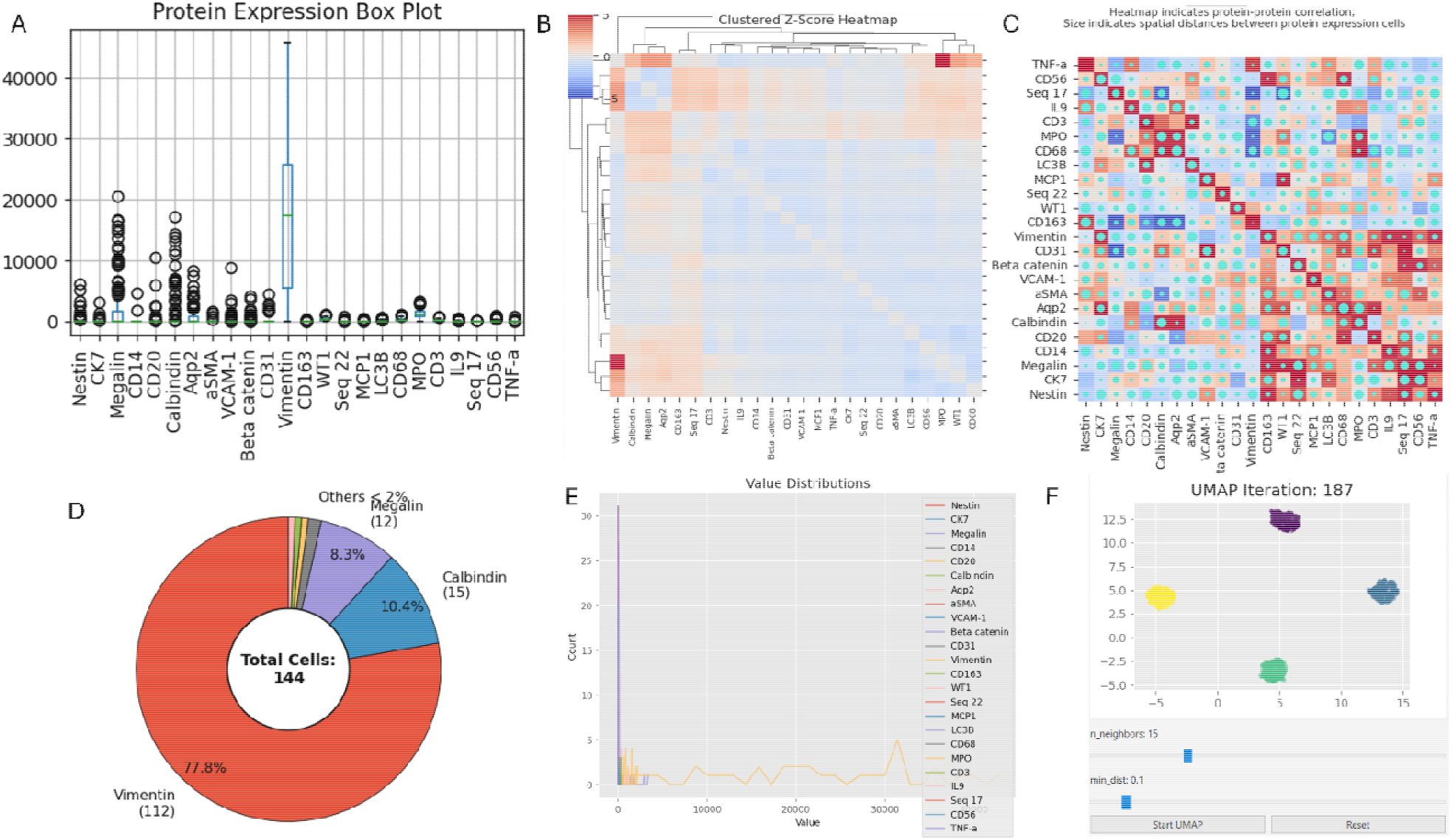
An sample output from MIST-Explorer for analyzing a kidney biopsy sample. A) Quantitative protein expression levels in single cells; B) Z-score clustering of proteins in single cells; C) Spatial proximity of protein-expressing cells; D) Cell composition in an region of interest; E) Distribution of protein expression values in cells; F) UMAP clustering of single cells.

ROI Data Filtering: Cell data corresponding to selected ROIs are extracted using precise geometric filtering algorithms: coordinate bounds checking for rectangles, elliptical equation evaluation for circles/ellipses, and a ray casting algorithm for polygons.

Protein Selection and ROI Navigation: Within each ROI, users can select specific subsets of proteins for analysis using a multi-select dropdown. An interface allows navigation (Back/Next buttons) between multiple defined ROIs for comparative analysis. Individual ROI analyses can be deleted.

Visualization Generation: Several types of plots are generated on-demand (“lazy loading”) for the data within the current ROI and the selected proteins, using the Matplotlib library.

Boxplots: Displaying the distribution (median, quartiles, outliers) of expression levels for selected proteins.

Heatmaps: Visualizing either Z-score normalized expression values across cells and selected proteins, or the spatial distribution of a chosen protein’s expression within the ROI.

Pie Charts: Showing the relative proportions of cells based on categorical features derived from protein expression (e.g., high/low expressors, if defined).

Histograms: Displaying the frequency distribution of expression values for selected proteins.

UMAP Plots: Generating Uniform Manifold Approximation and Projection plots for dimensionality reduction and visualization of cell populations based on the expression profiles of selected proteins within the ROI. Interactive Plot Handling: Generated plots can be saved as image files (e.g., PNG, SVG) or expanded into separate, resizable windows for detailed inspection and comparison.

## 3 Discussion

We have successfully developed the MIST-Explorer, an intuitive application designed to facilitate advanced analysis of multiplexed data. Although initially tailored for the Spatial MIST technology, the Explorer’s flexible framework allows it to be readily extended to other microarray-based assays employing similar registration-based analytical methods including those used in genomic and transcriptomic studies. By eliminating the necessity for prior bioinformatics expertise or coding skills, MIST-Explorer enables researchers to efficiently analyze complex data sets, allowing them to focus directly on scientific interpretation and discovery. Consequently, the MIST-Explorer significantly broadens access to sophisticated analytical tools within the multiplexing research community. Ultimately, this tool holds the potential to accelerate scientific discoveries by simplifying data analysis and enhancing interpretability of multiplexed datasets.

## AUTHOR INFORMATION

### Author Contributions

\# C.F and J.C. contribute equally to this work.

### Notes

JW is the founder and scientific advisor of MIST Bioscience LLC.

## ACKNOWLEDGEMENT

This work was supported by the National Institutes of Health R21DK138409 (JW) and R35GM151972 (JW).

